# Defining tissue-and disease-associated macrophages using a transcriptome-based classification model

**DOI:** 10.1101/532986

**Authors:** Hung-Jen Chen, Andrew Y.F. Li Yim, Guillermo R. Griffith, Wouter J. de Jonge, Marcel M.A.M. Mannens, Enrico Ferrero, Peter Henneman, Menno P.J. de Winther

**Affiliations:** Department of Medical Biochemistry, Experimental Vascular Biology, Amsterdam University Medical Centers, University of Amsterdam, Amsterdam, the Netherlands; Genome Diagnostics Laboratory, Department of Clinical Genetics, Amsterdam University Medical Centers, University of Amsterdam, Amsterdam, the Netherlands; Epigenetics Discovery Performance Unit, GlaxoSmithKline, Stevenage, United Kingdom; Tytgat Institute for Liver and Intestinal Research, Amsterdam University Medical Centers, University of Amsterdam, Amsterdam, the Netherlands; Institute for Cardiovascular Prevention (IPEK), Munich, Germany; Computational Biology, Target Sciences, GlaxoSmithKline, Stevenage, United Kingdom

## Abstract

Macrophages are heterogeneous multifunctional leukocytes which are regulated in a tissue-and disease-specific context. Many different studies have been published using *in vitro* macrophage models to study disease. Here, we aggregated public expression data to define consensus expression profiles for eight commonly-used *in vitro* macrophage models. Altogether, we observed well-known but also novel markers for different macrophage subtypes. Using these data we subsequently built the classifier macIDR, capable of distinguishing macrophage subsets with high accuracy (>0.95). This classifier was subsequently applied to transcriptional profiles of tissue-isolated and disease-associated macrophages to specifically define macrophage characteristics *in vivo*. Classification of these *in vivo* macrophages showed that alveolar macrophages displayed high resemblance to interleukin-10 activated macrophages, whereas macrophages from patients with chronic obstructive pulmonary disease patients displayed a drop in interferon-γ signature. Adipose tissue-derived macrophages were classified as unstimulated macrophages, but resembled LPS-activated macrophages more in diabetic-obese patients. Finally, rheumatoid arthritic synovial macrophages showed characteristics of both interleukin-10 or interferon-γ signatures. Altogether, our results suggest that macIDR is capable of identifying macrophage-specific changes as a result of tissue-and disease-specific stimuli and thereby can be used to better define and model populations of macrophages that contribute to disease.

## MAIN TEXT

### Introduction

Macrophages are multifunctional innate immune cells that play a central role in the spatiotemporal regulation of tissue homeostasis between pro-inflammatory defense and anti-inflammatory tissue repair. Dysregulation of macrophages has been implicated in a variety of disorders. As *in vivo* macrophages are often difficult to obtain and study, *in vitro* peripheral blood monocyte derived macrophages (MDMs) have been used extensively as model systems for assessing the transcriptional and functional regulation in response to various stimuli.

To mimic *in vivo* macrophages encountering different microenvironmental signals (*1, 2*), MDMs, differentiated with for example macrophage-(M-CSF), or granulocyte-macrophage stimulating factor (GM-CSF), are activated *in vitro* with bacterial lipopolysaccharides (LPS) or Th1 cytokine interferon gamma (IFNγ) to generate pro-inflammatory macrophages (M1). Anti-inflammatory macrophages (M2) are often generated by activating the cells with Th2 cytokines, interleukin-4 (IL4), or other anti-inflammatory stimuli, such as interleukin-10 (IL10), tumor growth factor (TGF) and glucocorticoids (*1-6*). By applying these pro-or anti-inflammatory stimuli to the MDMs, researchers sought to obtain proper models for studying the transcriptomic alteration associated with inflammation, infection, wound healing, and tumor growth. However, compared to *in vitro* MDM activation, *in vivo* macrophage activation represents a complex and dynamic process driven by multiple local factors. The most comprehensive study to date on gene expression profiling of *in vitro* macrophages was performed by Xue *et al.*, where the authors activated MDMs under various conditions and identified distinct stimulus-specific transcriptional modules (*5*). Their results suggested a broader view of how macrophages react upon divergent stimulation, thereby extending the classical dichotomous pro-and anti-inflammatory model to an activation spectrum. While there have been attempts to summarize published studies in an effort to attain consensus (*4*), a proper integrative analysis has thus far not been performed. Furthermore, it remains unclear to what extent the *in vitro* macrophage-models resemble their *in vivo* counterparts and which specific subtypes associate with disease.

In this study, we integrated 206 microarray and RNA-sequencing (RNA-seq) datasets from 19 different studies (*7-24*) (Table 1) to systematically characterize eight different human MDM activation states. First, we identified consistently differentially expressed genes (cDEGs) by comparing activated with unstimulated macrophages using a random effects meta-analysis (*25*). Next, we implemented penalized multinomial logistic regression to construct a classification model, macIDR (<u>mac</u>rophage <u>id</u>entifier), capable of distinguishing specific MDM activation states, independent of the macrophage differentiation factor used. Finally, we used macIDR to project *in vivo* tissue-and disease-associated macrophages onto the eight *in vitro* MDMs (*26-34*) and to identify expression signatures derived from the tissues and specific for the patient groups.

**Table 1.** Included datasets. An overview of the datasets and the associated studies included in the in the meta-analysis and the classification analysis.

| Reference | Type | Dataset ID | Purpose |
| --- | --- | --- | --- |
| Fuentes-Duculan et al. 2010 | Microarray | GSE18686 | Meta-analysis, training & test |
| Schroder et al. 2012 | Microarray | GSE19765 | Meta-analysis, training & test |
| Benoit et al. 2012 | Microarray | GSE30177 | Meta-analysis, training & test |
| Chandriani et al. 2014 | Microarray | GSE47538 | Meta-analysis, training & test |
| Martinez et al. 2005 | Microarray | GSE5099* | Meta-analysis, training & test |
| Derlindati et al. 2014 | Microarray | GSE57614 | Meta-analysis, training & test |
| Jubb et al. 2016 | Microarray | GSE61880 | Meta-analysis, training & test |
| Steiger et al. 2016 | Microarray | GSE79077 | Meta-analysis, training & test |
| Fujiwara et al. 2016 | Microarray | GSE85346 | Meta-analysis, training & test |
| Corbi et al. 2017 | Microarray | GSE99056 | Meta-analysis, training & test |
| Tsang et al. 2009 | Microarray | E-MEXP-2032 | Meta-analysis, training & test |
| Przybyl et al. 2016 | Microarray | E-MTAB-3309 | Meta-analysis, training & test |
| Turner et al. 2017 | Microarray | E-MTAB-5095 | Meta-analysis, training & test |
| Surdziel et al. 2017 | Microarray | E-MTAB-5913 | Meta-analysis, training & test |
|  | RNAseq | BLUEPRINT | Meta-analysis, training & test |
| Park et al. 2017 | RNAseq | GSE100382 | Meta-analysis, training & test |
| Zhang et al. 2015 | RNAseq | GSE55536 | Meta-analysis, training & test |
| Martins et al. 2017 | RNAseq | GSE80727 | Meta-analysis, training & test |
| Realegeno et al. 2016 | RNAseq | GSE82227 | Meta-analysis, training & test |
| Own | RNAseq | E-MTAB-7572 | Test |
| Riera-Borull et al. 2017 | Microarray | GSE99056 | GM-CSF verification |
| Vento-Tormo et al. 2016 | Microarray | GSE75938 | GM-CSF verification, non-MDM |
| Xue et al. 2014 | Microarray | GSE46903 | GM-CSF verification, non-MDM |
| Tasaki et al. 2018 | Microarray | GSE93776 | GM-CSF verification, non-MDM |
| Yarilina et al. 2008 | Microarray | GSE10500 | Synovial macrophage test |
| You et al. 2014 | Microarray | GSE49604 | GM-CSF verification, Synovial macrophage test |
| Shaykhiev et al. 2009 | Microarray | GSE13896 | Alveolar macrophage test |
| Woodruff et al. 2005 | Microarray | GSE2125 | Alveolar macrophage test |
| Madore et al. 2010 | Microarray | GSE22528 | Alveolar macrophage test |
| Goleva et al. 2008 | Microarray | GSE7368 | Alveolar macrophage test |
| Dalmas et al. 2014 | Microarray | GSE54350 | Adipose tissue macrophage test |
| Kang et al. 2017 | Microarray | GSE97779 | Synovial macrophage test |
| Asquith et al. 2013 | Microarray | E-MEXP-3890 | Synovial macrophage test |

\*The GSE5099 dataset was composed of two Affymetrix microarray datasets: U133A and U133B. Due to the limited overlap in genes between U133B with the rest, we only included the U133A dataset.

### Results

#### Meta-analysis defines both well-known and novel transcriptional markers of macrophage activation states

We searched the public repositories Gene Expression Omnibus and ArrayExpress for MDMs differentiated with macrophage colony-stimulating-factor (M-CSF) and activated with commonly used stimuli. To be included in the analysis, each activated MDM alongside an unstimulated control had to be represented by at least 4 different studies, with each study containing at least 2 biological replicates per activation state. In total, we assembled a cohort for eight macrophage activation states: unstimulated macrophages (M0) and macrophages activated by: short exposure (2 to 4 hours) to LPS (M-LPS_early_) or long exposure (18 to 24 hours) to LPS (M-LPS_late_), LPS with IFNγ (M-LPS+IFNγ), IFNγ (M-IFNγ), IL-4 (M-IL4), IL-10 (M-IL10), and dexamethasone (M-dex) (Table 1). To find consistent transcriptional differences, we performed a random effects meta-analysis where we calculated the standardized effect size per study by comparing each macrophage activation state with M0.

We identified consistent differentially expressed genes (cDEGs) that were previously observed to be characteristic for certain *in vitro* macrophage subsets (Table S1). For example, interleukin-1 beta (*IL1B*), C-C Motif Chemokine Ligand 17 (*CCL17*) and Cluster of Differentiation 163 (*CD163*) were consistently upregulated when comparing M-LPS_late,_ M-IL4, and M-dex with M0, respectively (*4*). Notably, we also identified several novel genes that were not widely considered as activated macrophage markers. Consistent upregulation of interleukin 7 receptor (*IL7R*) and CD163 Molecule Like 1 (*CD163L1*) was observed when comparing M-LPS_early_ and M-IL10 with M0, respectively. On the other hand, consistent downregulation of Nephroblastoma Overexpressed (*NOV*, also known as *CCN3*) and Adenosine A2b receptor (*ADORA2B*) was observed when comparing M-IFNγ, M-LPS_late_, and M-LPS+IFNγ with M0, respectively.

#### Correlation and enrichment analyses display classical pro-and anti-inflammatory clustering

Pairwise correlation analysis of the standardized effect sizes across studies showed that M-LPS_early_, M-LPS_late_, M-IFNγ, and M-LPS+IFNγ formed one cluster, whereas M-IL4, M-IL10 and M-dex formed a second cluster (Fig. 1A). Other activation states could not be discerned easily, which was attributed to study-specific effects. This observation agrees with previous studies where transcriptional alterations induced with these conventional pro-and anti-inflammatory stimuli were found to cluster according to the M1-M2 dichotomy (*2, 5*). Further sub-clustering appeared to divide the macrophages according to the stimuli. The separation between the M1 and the M2 was also apparent as the first principal component (PC1), which displayed a clear separation between M1 and M2 associated stimuli on the left and right, respectively (Fig. 1B). Relative to the M1 subsets, the M2 subsets appeared to display a stronger diversity along PC2, suggesting a more distinctive transcriptional programming of the individual subsets.

**Fig. 1.**
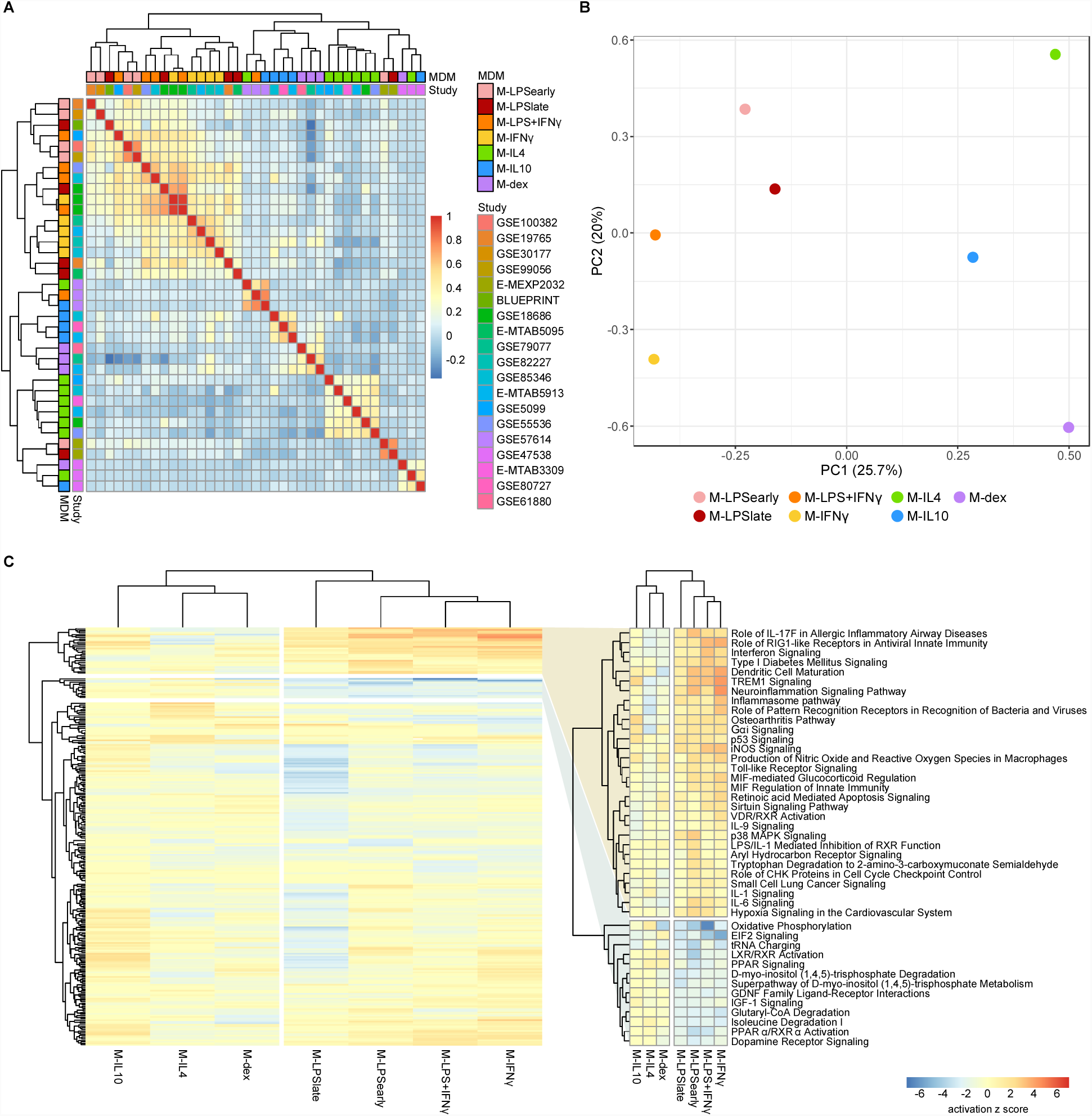
Summary meta-analysis. **(A)** Heatmap of the Cohen d pairwise Spearman correlation coefficients. **(B)** Principal component analysis of the Z-values obtained from the meta-analysis. **(C)** Heatmap of the canonical pathways with the intensity representing the activation z score. Two most defining clusters have been enlarged and annotated on the right.

Next, we performed canonical pathway analysis using the Ingenuity Pathway Analysis (IPA) software package. While the clusters were slightly different among individual macrophage subsets, the overall separation between M1 and M2 was still apparent with hierarchical clustering revealing two sets of pathways that appeared to be responsible for the separation (Fig. 1C and Table S1). Pathways known for their pro-and anti-inflammatory responses, such as interferon signaling and LXR/RXR activation, displayed a clear and distinct pattern for the M1 and M2 subsets, respectively.

#### A multinomial elastic net classifier distinguishes macrophage activation states with high accuracy

Subsequently, we sought to perform feature selection to identify genes capable of distinguishing macrophage activation states. We merged the data from different microarray and RNA-seq platforms and performed multinomial elastic net regression on the 5986 overlapping genes present in all datasets. The expression data was randomly split into a training (2/3) and a test (1/3) set whereupon the training set was used to build a model through ten-fold cross-validation. We repeated the cross-validation procedure 500 times and took the median thereof to stabilize the log odds ratios (*35*). Genes were considered stable predictors if their log odds ratio was non-zero in more than 50% of the iterations (Fig. S1). Subsequent classification was performed using the median log odds ratio per gene across all iterations (Fig. 2A). Altogether, our classifier was composed of 97 median-stabilized predictor genes, and was compiled as an R package called macIDR (https://github.com/ND91/macIDR).

**Fig. 2.**
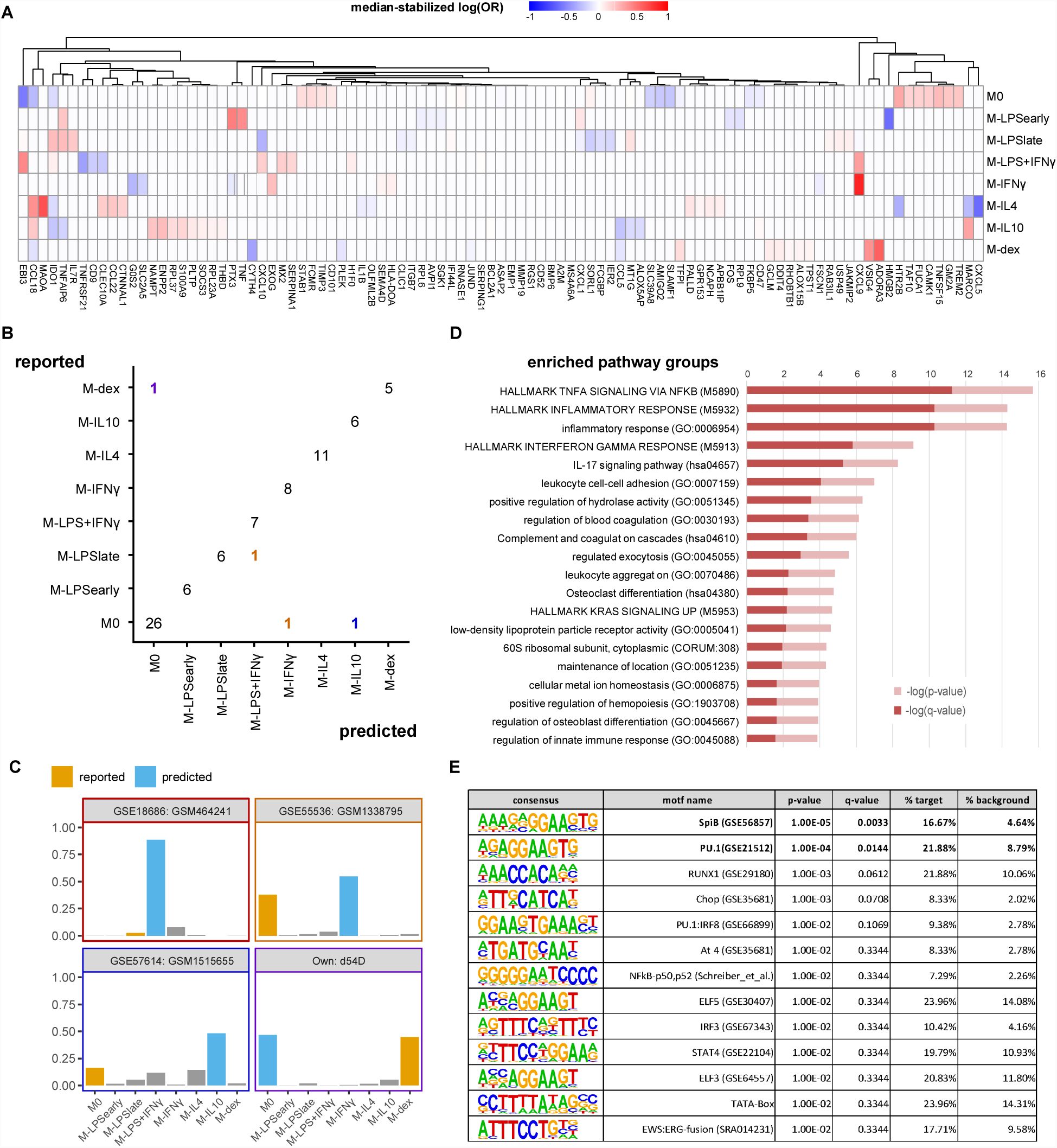
Classification results. **(A)** Heatmap of the median-stabilized log odds ratios per macrophage activation state for each of the 97 predictor genes. **(B)** Confusion matrix representing the number of correctly classified samples (entries on the diagonal) versus the misclassified samples (entries on the off-diagonal). Classes on the y-axis represents the reported class while classes on the x-axis represent the predicted class. **(C)** Bar plots of the misclassified samples depicting the classification signal on a scale of 0 to 1 where the class with the largest signal represents the predicted class. Blue bars represent the incorrectly predicted class and orange bars represent the reported class. **(D)** Pathway overrepresentation analysis of the predictor genes ranked by p-value. **(E)** Motif overrepresentation analysis of the predictor genes ranked by *p*-value. Columns represent the consensus sequence, the motif name, the *p*-value, the BH-adjusted *p*-value (q-value), the percentage of provided genes with the motif, and the percentage genes in the background with the motif.

To validate our model, we tested macIDR against the previously withheld test set, which included a newly-generated RNA-seq experiment containing all included activation states. Classification of the test set revealed an accuracy above 0.95 with both high sensitivity (>0.98) and specificity (> 0.83; Table 2). In total, 75 out of 79 test samples were correctly classified (Fig. 2B). Notably, for three of the four misclassified samples, the second-best prediction was the subset as reported by the authors. Investigation of the four misclassified samples revealed that most errors were made regarding M0: two M0 datasets were classified as M-IL10 (GSM151655) and M-IFNγ (GSM1338795), and one M-dex dataset (d54D) as M0. The fourth misclassification pertained a M-LPS_late_ (GSM464241) dataset classified as M-LPS+IFNγ (Fig. 2C).

**Table 2.** Classification testing. A confusion matrix representing the classifier performance on the test set. TP: True positives, FP: False positives, TN: True negatives, FN: False negatives, TNR: True negative rate/specificity, TPR: True positive rate/sensitivity.

| Macrophage | TP | FP | TN | FN | TNR | TPR | Accuracy |
| --- | --- | --- | --- | --- | --- | --- | --- |
| **M0** | 26 | 1 | 50 | 2 | **0.98** | **0.93** | **0.96** |
| **M-LPS<sub>early</sub>** | 6 | 0 | 73 | 0 | **1.00** | **1.00** | **1.00** |
| **M-LPS<sub>late</sub>** | 6 | 0 | 72 | 1 | **1.00** | **0.86** | **0.99** |
| **M-LPS+IFNγ** | 7 | 1 | 71 | 0 | **0.99** | **1.00** | **0.99** |
| **M-IFNγ** | 8 | 1 | 70 | 0 | **0.99** | **1.00** | **0.99** |
| **M-IL4** | 11 | 0 | 68 | 0 | **1.00** | **1.00** | **1.00** |
| **M-IL10** | 6 | 1 | 72 | 0 | **0.99** | **1.00** | **0.99** |
| **M-dex** | 5 | 0 | 73 | 1 | **1.00** | **0.83** | **0.99** |

Pathway analysis of the predictor genes revealed a clear enrichment for inflammatory pathways, such as TNFα signaling, inflammatory response and interferon gamma signaling, confirming the importance of inflammation regulation in macrophage activation (Fig. 2D, Table S3). A follow-up transcription factor motif analysis on the promoters of the predictor genes showed significant enrichment for macrophage transcription factors, the E26 transformation-specific PU.1 (Spi1) and SpiB (Fig. 2E).

#### MacIDR generalizes to granulocyte-macrophage colony-stimulating-factor differentiated macrophages

Besides M-CSF, granulocyte-macrophage colony-stimulating-factor (GM-CSF) is often used to differentiate monocytes to macrophages, where it is thought to evoke a more pro-inflammatory phenotype (*36-38*). We investigated whether macIDR was capable of discerning GM-CSF-differentiated MDMs (GM-MDMs) exposed to various stimuli. In total, we obtained 31 datasets from three microarray studies (*5, 16, 39*) where GM-MDMs were activated with LPS (GM-LPS_early_, GM-LPS_late_), IFNγ (GM-IFNγ), and IL4 (GM-IL4), or remained unstimulated (GM0). These studies were selected based on their similarity in activation duration with the M-CSF differentiated macrophages used in the training set. We observed that all GM-MDMs were classified as their M-CSF counterparts (Fig. 3B) despite the fact that some predictive genes were absent from two studies (Table S4). Our results indicate that the predictor genes for M-LPS_early_, M-LPS_late_, M-IFNγ and M-IL4 are representative for the activation regardless of whether M-CSF or GM-CSF was used for differentiation.

**Fig. 3.**
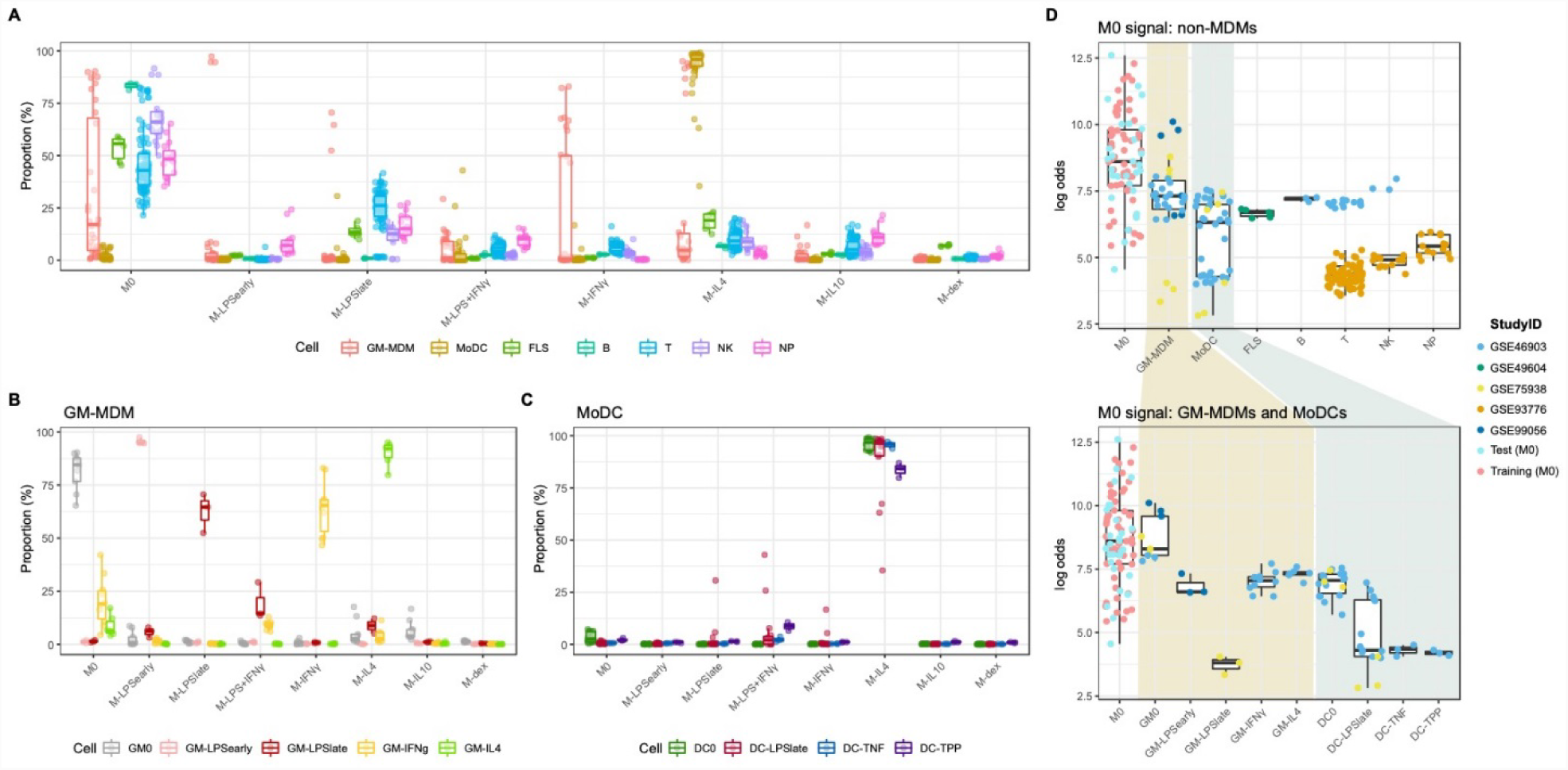
Classification of cells not included in training set. **(A)** Boxplots representing the classification signal on a scale of 0 to 1 where classes with the largest signal represents the predictions. Colors represent GM-CSF differentiated macrophages (GM-MDMs), monocyte-derived dendritic cells (MoDCs), fibroblast-like synoviocytes (FLS), B lymphocytes (B), T lymphocytes (T), natural killer cells (NK), and neutrophils (NP). **(B)** GM-MDMs and **(C)** MoDCs colored by stimulation. **(D)** Visualization of the log odds distribution of the M0 signal for the M0s from training and test set and the non-macrophage immune cells. Further separation across different stimulations for the GM-MDMs and the MoDCs was plotted below.

#### Non-macrophage cells classify as M0 whereas monocyte-derived dendritic cells classify as M-IL4

To understand the limitations of macIDR, we investigated its performance on non-MDM cells. Datasets were obtained of monocyte-derived dendritic cells (MoDC) (*5, 39*), various T lymphocyte subtypes (T), B lymphocytes (B), neutrophils (NP), and natural killer cells (NK) (*16, 40*), as well as fibroblast-like synoviocytes (FLS) (*27*).

The MoDCs were primarily recognized as M-IL4 regardless of subsequent LPS maturation or other pro-inflammatory stimulation (Fig. 3C). Spurious classification was ruled out as GM0 and GM-LPS_late_ datasets from the same studies were correctly classified as described previously. Investigation of the M-IL4 predictor genes revealed that concordant expression of monoamine oxidase A (*MAOA*) and lower discordant expression of C-X-C motif chemokine 5 (*CXCL5*) of MoDCs were likely the main contributing factors towards the M-IL4 classification (Table S4). By contrast, we found that the FLS, T, B, NP and NK cells were all classified as M0 (Fig. 3A), suggesting that the M0 class is used as a label for expression profiles of non-monocyte-derived cells. By looking at the distribution of the log odds we observed that true M0 classifications generally displayed higher log odds than the M0 classifications of most non-macrophage cells, but found this not to be definite (Fig. 3D). The increased variance for the M0 signal observed for GM-MDMs and MoDCs was due to the different GM-MDMs activation and DC maturation states, as GM0 and up to some extent DC0, depicted M0 log odds similar to true M0 samples.

#### Alveolar macrophages from COPD and smoking individuals show a reduced M-IFNγ signal

We next attempted to classify macrophages derived from patient tissues to study their semblance to *in vitro* generated MDMs. We investigated alveolar macrophages (AMs) obtained through bronchioalveolar lavage from smoking individuals, COPD patients, asthma patients (*31*), and healthy control (HCs). Overall, we found that the AMs were classified primarily as M-IL10 (Fig. 4A and Table S10), which appears to be driven by the *MARCO* signal. This corroborates the observation that lung tissue, specifically AMs, display significantly higher gene expression of *MARCO* relative to surrounding cells and other tissues (*41, 42*). Moreover, *MARCO* was found to be necessary in AMs for mounting a proper defense response (*41-43*). Further comparisons of the AMs derived from different patient groups, we found that the macrophage signal could be stratified according to health status where COPD-and smoker-derived AMs displayed a higher M-IL4, M-IL10 and M-dex signal and a reduced M-IFNγ and M-LPS+IFNγ signal compared to AMs obtained from HCs. This observation indicates a stronger M2-and lesser M1-like phenotype, which was also noted in previous studies (*5, 31*). This difference in classification signal was found to be driven by a decreased log odds for *CXCL9* (M-IFNγ and M-LPS+IFNγ) and *CXCL5* (M-IL4) and increased log odds for *TNFAIP6* (M-IL10), and *ADORA3* (M-dex; Fig. S2).

**Fig. 4.**
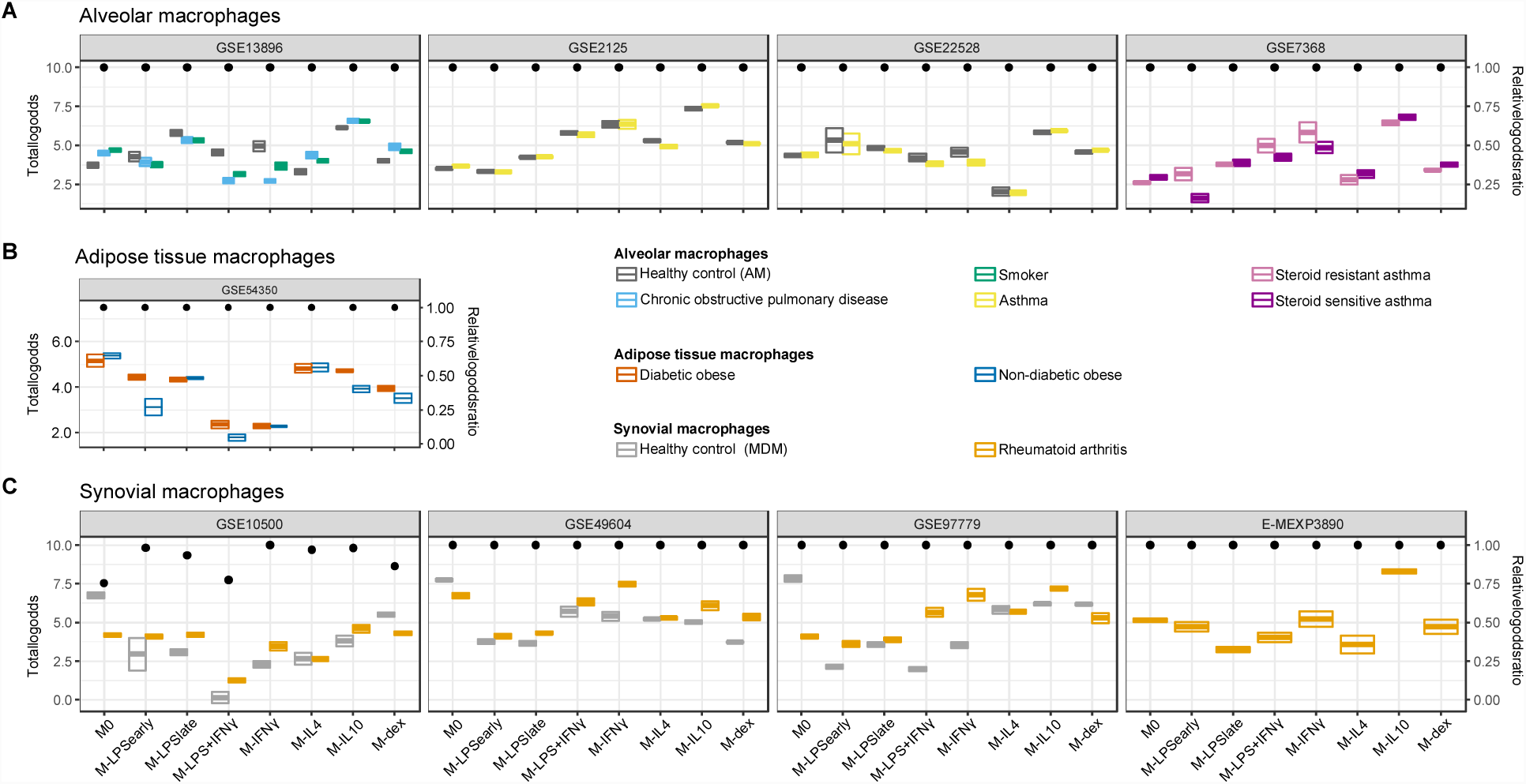
Classification of in vivo macrophages. Summarized classification results per dataset with cross-bars representing the mean and the standard errors of the log odds colored by the macrophage in vivo type. Dots above represent the log odds ratio (log(OR)) relative to the sum of the log odds ratios if all predictor genes were measured. **(A)** Alveolar macrophages obtained from smoking individuals, chronic obstructive pulmonary disease (COPD), asthma patients, as well as healthy controls. **(B)** Adipose tissue macrophages obtained from diabetic obese and non-diabetic obese patients. **(C)** Synovial macrophages obtained from rheumatoid arthritis (RA) patients and MDMs from healthy controls (HCs).

Furthermore, while no clear differences were observed when comparing AMs from asthma patients to HCs, AMs from steroid-sensitive asthma patients displayed a decreased M-dex signal and increased M-LPS_early_, M-IFNγ and M-LPS+IFNγ signal relative to steroid resistant asthma AMs (Fig. 4A). The apparent difference in steroid sensitivity appeared to be caused by a difference in *TNF* and *CXCL9* signals contributing to the M-IFNγ and M-LPS+IFNγ classification respectively (Fig. S2). This observation corroborates previous studies where IFNγ signaling was found to suppress glucocorticoid-triggered transcriptional remodeling in macrophages leading to the macrophage-dependent steroid-resistance, thereby reflecting a higher level of IFNγ in steroid-resistant relative to steroid-sensitive asthma patients.

#### Adipose macrophages show a M0 and M-IL4 classification

We next investigated visceral adipose tissue macrophages (ATM) derived from diabetic and non-diabetic obese patients (*34*). Classification analysis suggested visceral ATMs showed most similarity with M0 followed by M-IL4 (Fig. 4B). While we were not capable of defining a set of genes responsible for the M0 classification, we observed that the concordant expression of *MAOA* and C-C chemokine ligand 18 (*CCL18*) contributed the most to the M-IL4 signal (Fig. S3). *MAOA* encodes a norepinephrine degradation enzyme and is expressed more in sympathetic neuron-associated macrophages isolated from the subcutaneous adipose tissue. In these cells, MAOA’s norepinephrine clearance activity has been linked to obesity (*44*). Interestingly, when comparing ATMs from diabetic obese with non-diabetic obese patients, we observed a stronger signal of M-IL10 and M-LPS_early_ driven by *CCL18* and *TNF* respectively (Fig. S3). Notably, *CCL18* expression in both visceral and subcutaneous adipose tissue has been associated with insulin-resistant obesity (*45, 46*). Furthermore, several studies have demonstrated that ATMs could be divided into CD11C^+^CD206^+^ and CD11C^-^CD206^+^ subpopulations (*47, 48*). Specifically, an increased density of the IL10 and TNFα-secreting CD11C^+^CD206^+^ ATMs in adipose tissue was associated with insulin resistance (*47*), which coincides with the observed increase of M-IL10 and M-LPS_early_ signals while the M-IL4 signal remained unaltered. Altogether, this observation shows the diverse roles of macrophages in obesity and highlights the complex crosstalk between neural signaling, immune system and metabolism.

#### Synovial macrophages from rheumatoid arthritis patients display similarity to M-IFNγ and M-IL10

Finally, we analyzed synovial macrophages (SMs) from RA patients and MDMs from HCs (*26-29*). Whereas unstimulated MDMs were successfully classified as M0, SMs from RA patients were classified as either M-IL10 or M-IFNγ (Fig. 4C). Specifically, RA-derived synovial macrophages (RA-SMs) from 3 studies (*26, 28, 29*) were classified as M-IL10, whereas samples from one study (*27*) were classified as M-IFNγ. Comparison of the classification signal of the RA-SMs with the HC MDMs displayed a higher signal for M-IFNγ and M-IL10 (Fig. 4C). Concordantly, a previous study reported an increased gene expression of *IFNG* and *IL10* in RA synovial fluid mononuclear cells compared with PBMCs from both RA patients and HCs (*49*). Further investigation of the M-IFNγ and M-IL10 classification revealed dominant signals for M-IFNγ predictor gene C-X-C Motif Chemokine Ligand 9 (*CXCL9*) and for M-IL10 predictor gene Macrophage Receptor With Collagenous Structure (*MARCO*; Fig. S4). This observation agrees with previous studies where elevated gene and protein expression of CXCL9 was found in the synovium of RA patients compared with that of osteoarthritis patients (*50, 51*). Similarly, an increased presence of MARCO was detected in the inflamed joints, particularly in RA patients (*52*).

### Discussion

In this study, we performed a macrophage characterization study by integrating public datasets of eight *in vitro* macrophage activation states. Our meta-analysis returned both well-known and novel markers for activated macrophages. At a genome-wide level, we observed separation according to the conventional pro-and anti-inflammatory macrophages. We subsequently built a classification model capable of discriminating macrophage activation states based on their transcriptomic profile and made this available as an R package called macIDR. By applying macIDR to *in vivo* macrophages, we projected the latter onto the eight *in vitro* macrophage models providing insights in how disease and tissue of origin affected the predicted composition.

Previous macrophage characterization studies focused primarily on gathering large cohorts. Xue *et al.* adopted an inclusive strategy by categorizing genes into different activation states through self-organizing maps and correlation analyses (5). Instead, we sought to find consensus from published data by implementing a descriptive and an exclusive strategy representing the meta-analysis and the elastic net classification analysis respectively. Where the meta-analysis identified genes that were consistently differentially expressed across studies when comparing stimulated with unstimulated MDMs, elastic net classification analysis represented a rigorous feature selection approach that yielded predictor genes capable of classifying the eight *in vitro* MDMs with high accuracy. We implemented an additional layer of robustness by performing repeated cross-validation to ensure that the final output of the elastic net regression was stable.

Many of the observed cDEGs and predictor genes have been recognized as bona fide markers for different activation states, such as *TNF* (M-LPS_early_), *IDO1* (M-LPS+IFNγ), *CXCL9* (M-IFNγ), and *ADORA3* (M-dex) (*4*) (Table S1 and Fig 2A). Pathway and transcription factor motif analyses of the predictor genes revealed enrichment for inflammatory pathways and macrophage transcription factors PU.1 and SpiB. As both PU.1 and SpiB are key transcription factors that drive macrophage differentiation (*53*), our observation suggests that activation is determined by the regulation of macrophage-specific inflammation.

To test the limits of the classification model, we classified non-MDM cells. Classification of MoDCs indicated that they were most similar to M-IL4, even after treatment with pro-inflammatory agents such as LPS. This observation is likely due to the method to generate MoDCs, where monocytes are differentiated with GM-CSF and IL4. Notably, while the MoDCs were classified primarily as M-IL4, some MoDCs matured with LPS for 24 hours displayed a mildly increased signal towards M-LPS+IFNγ and M-LPS_late_. Similarly, immature MoDCs (DC0) displayed a slightly higher response for M0. All other non-macrophage cells were classified as M0. While the log odds of proper M0 classifications were slightly higher than improper M0 classifications, we acknowledge that no clear threshold could be defined. We are unsure why M0 was predicted as a class for non-MDM and non-MoDC datasets, but we speculate that it might be related to the M0 class being represented by most predictor genes relative to the other classes. We therefore recommend potential users of macIDR to determine *a priori* that their dataset of interest represents macrophages either experimentally or using *in silico* cell-composition estimation methods, such as CIBERSORT (*54*) or xCell (*55*).

As macIDR was capable of properly classifying differentially differentiated *in vitro* MDMs, we applied it to *in vivo* macrophages with the goal of extracting subpopulation information. When comparing tissue-derived macrophages among different patient groups and healthy donors, we observed differences in predictions, such as reduced signals of M-IFNγ and M-LPS+IFNγ and increased signals of M-IL4, M-IL10 and M-dex when comparing AMs from COPD patients with HCs. Moreover, we observed that the tissue of origin had a large impact on the macrophage classification with *in vivo* macrophages obtained from HCs not always being classified as M0 as was observed for the AMs. Since some *in vivo* macrophages obtained from healthy tissue were classified as *in vitro* activated MDMs, unstimulated *in vitro* MDMs likely do not reflect the basal state of all tissue-resident macrophages underpinning the importance of how multiple factors in the microenvironment shape the transcriptome. Our results suggest that for some *in vitro* models, using activated MDMs might achieve a more comparable phenotype to the *in vivo* tissue macrophages that express the tissue transcriptomic signatures.

Unlike the AMs, no transcriptomic data was available of SMs from healthy donors. We were therefore unable to conclude whether the M-IL10 and M-IFNγ predictions observed for the samples from RA patients were tissue-specific or disease-associated. Though samples from different studies were classified as M-IL10 or M-IFNγ, these two activation states appeared to be the highest two predicted classes for SMs from RA patients among all recruited datasets. Notably, M-IL10 and M-IFNγ represent predictions on opposite sides of the conventional inflammatory spectrum. Characterization studies on RA-SMs suggested that they represent multiple subpopulations, such as the CD163^+^ anti-inflammatory tissue-resident macrophages and the S100A8/9^**+**^ **pro-**inflammatory macrophages recruited from peripheral monocytes (*56*).

As *in vivo* macrophages likely represent a more heterogeneous population compared to the *in vitro* MDMs used in building the classifier, prediction of macrophage mixtures using bulk RNA-seq or microarray returns signals from multiple different subsets. The classification therefore mainly reflects altered subset proportions due to disease progression or association. It is likely that some *in vivo* macrophage subpopulations do not share transcriptomic signatures with any of the eight *in vitro* MDMs preventing the exact characterization thereof. Future studies using single cell RNA-sequencing should aim at defining novel additional *in vivo* macrophage subsets. Nonetheless, we were capable of extracting signals from the *in vivo* macrophage classifications that not only corroborated previous findings, but also provided novel features for future research. It is essential to analyze macrophage subsets within the context of their *in vivo* environment and therefore we provide this quantitative method to aid researchers in better defining and modelling macrophages in tissue and disease.

### Materials and Methods

#### Data selection

Datasets were found through an extensive search of the National Center for Biotechnology Information (NCBI) Gene Expression Omnibus (GEO) and European Bioinformatics Institute (EBI) ArrayExpress (AE). For our search, we used the keywords “(macrophage) OR (monocyte) OR (MDM) OR (HBDM) OR (MoDM) OR (MAC) OR (dendritic cell) AND “Homo sapiens” [porgn:__txid9606]”. Our search yielded 1,851 and 175 experiments for GEO and AE respectively at the time of writing (May 2018).

The initial screen was limited to studies that investigated primary macrophages and excluded the stem cell derived macrophages or immortalized cell lines. Then we categorized macrophage subsets based on the stimuli and the treatment time. For each subset, we sought to obtain at least 4 studies including at least 2 biological replicates. Further, as a background control, only studies including unstimulated control macrophages were selected. After this screening, we investigated macrophages stimulated with control medium, LPS (either for 2 to 4 hours or 18 to 24 hours), LPS with IFNγ, IFNγ, IL4, IL10 or dexamethasone for 18 to 24 hours. Microarray datasets generated on platforms other than Illumina, Affymetrix or Agilent were excluded to ensure comparability. As we only investigated genes that were measured in every single study, datasets that displayed limited overlap in the measured genes with the other studies were removed. The *in vivo* macrophage datasets were samples obtained from clinical specimen. In total, we obtained 206 datasets belonging to 19 studies for the meta-analysis and classification.

#### Meta-analysis

A random effects meta-analysis was performed on the normalized data using the GeneMeta (v1.52.0) (*57*) package, which implements the statistical framework outlined by Choi *et al.* (*25*). In short, the standardized effect size (Cohen d adjusted using Hedges and Olkin’s bias factor) and the associated variance were calculated for each study by comparing each activated macrophage with the unstimulated macrophage within each study. The standardized effect sizes were then compared across studies by means of random effects model to correct for the inter-study variation, thereby yielding a weighted least squares estimator of the effect size and its associated variance. The estimator of the effect size was then used to calculate the Z-statistic and the *p*-value, which was corrected for multiple testing using the Benjamini-Hochberg procedure. We modified the GeneMeta functions to incorporate the shrunken sample variances obtained from limma (*58*) for calculating the standardized effect sizes.

#### Classification

Raw microarray and RNA-seq data were log_2_ transformed where necessary after which the data was (inner) merged and randomly divided into a training (2/3) and test set (1/3). As the test set should remain hidden from the training set, raw microarray and RNA-seq data was used instead of normalized data to prevent data leakage (*59*). Elastic net regression was subsequently performed using the R glmnet (v2.0) (*60*) package. As penalized regression approaches are sensitive to the magnitude of each feature, we investigated the use of standardization. As the lowest deviance appeared to be higher in the inter-fold standardized training set relative to raw data, we did not include any standardization (Fig. S1). We also investigated the optimal alpha for minimizing the deviance by means of an initial grid search approach, which was found to be 0.8.

The training set was subjected to ten-fold cross-validation for tuning the penalty regularization penalty parameter lambda. This process was subsequently repeated 500 times to stabilize the randomness introduced during the splitting step for cross-validation (*35*). We considered genes to be stable classifiers if they displayed a non-zero log odds ratio in at least 50% of the 500 iterations. The final log odds ratio was selected by taking the median of each stable predictor gene across the 500 iterations. We subsequently validated our classifier on the withheld test set.

In addition to the training and test data, we downloaded and imported additional datasets from GM-MDMs, non-macrophage cells, and *in vivo* macrophages. Subsequent classification was performed using the macIDR package. Unlike the studies included for training and testing, some of the included studies were performed on platforms that did not measure the expression of some of the predictor genes. To that end, the relative log odds ratios were calculated, which represent the log odds ratio present relative to the total log odds ratio had all predictor genes been present.

#### Human monocyte-derived macrophage differentiation and stimulation

Buffy coats from three healthy anonymous donors were acquired from the Sanquin blood bank in Amsterdam, the Netherlands. Monocytes were isolated through density centrifugation using Lymphoprep(tm) (Axis-Shield) followed by human CD14 magnetic beads purifcation with the MACS^®^ cell separation columns (Miltenyi). The resulting monocytes were seeded on 24-well tissue culture plates at a density of 0.8 million cells/well. Cells were subsequently differentiated to macrophages for 6 days in the presence of 50ng/mL human M-CSF (Miltenyi) with Iscove’s Modified Dulbecco’s Medium (IMDM) containing 10% heat-inactivated fetal bovine serum, 1% Penicillin/Streptomycin solution (Gibco) and 1% L-glutamine solution (Gibco). The medium was renewed on the third day. After differentiation, the medium was replaced by culture medium without M-CSF and supplemented with the following stimuli: nothing, 10ng/mL LPS (Sigma, *E. coli* E55:O5), 10ng/mL LPS plus 50ng/mL IFNγ (R&D), 50ng/mL IFNγ, 50ng/mL IL4 (PeproTech), 50ng/mL IL10 (R&D), 100nM dexamethasone (Sigma) for 24 hours. LPS_early_ macrophages were first cultured with culture medium for 21 hours and then stimulated with 10ng/mL LPS for 3 hours prior to harvest.

#### RNA isolation and sequencing library preparation and analyses

Total RNA was isolated with Qiagen RNeasy Mini Kit per the manufacturer’s recommended protocol. RNA sequencing libraries were prepared using the standard protocols of NuGEN Ovation RNA-Seq System V2 kit. Size-selected cDNA library samples were sequenced on a HiSeq 4000 sequencer (Illumina) to a depth of 16M per sample according to the 50 bp single-end protocol at the Amsterdam University Medical Centers, location Vrije Universiteit medical center. Raw fastq files were subsequently processed in the same manner as the public datasets to maintain consistency.

## Supporting information

Table S1 meta-analysis results

Table S2 meta-analysis pathway analysis

Table S3 predictor gene pathway analysis

Table S4 Log odds for the verification datasets

## General

We are thankful to all authors and participants of the studies included in this research for sharing their transcriptomic data. This study makes use of data generated by the Blueprint Consortium. A full list of the investigators who contributed to the generation of the data is available from www.blueprint-epigenome.eu. Funding for the project was provided by the European Union’s Seventh Framework Programme (FP7/2007-2013) under grant agreement no 282510 – BLUEPRINT

## Author contributions

HJC performed the literature search. HJC and GG cultured the cells, isolated the RNA for the RNA sequencing analysis. AYFLY downloaded, cleaned and performed the *in silico* analyses of the public and in-house generated data. HJC, AYFLY, PH and MPJW decided on the inclusion criteria for the public data. HJC and AYFLY drafted the manuscript. PH and MPJW conceived the study and participated together with EF, WJJ and MMAMM in the design and coordination of the study. All authors read and approved the final manuscript.

## Data and materials availability

The RNA-seq data generated and analyzed during the current study are available in the ArrayExpress repository E-MTAB-7572. All public datasets analyzed during the current study are available in the Gene Expression Omnibus and ArrayExpress repositories. Studies from the GEO repository include: GSE18686, GSE19765, GSE30177, GSE47538, GSE5099, GSE57614, GSE61880, GSE79077, GSE85346, GSE99056, GSE100382, GSE55536, GSE80727, GSE82227, GSE99056, GSE75938, GSE46903, GSE93776, GSE10500, GSE49604, GSE97779, and GSE13896. Studies from the ArrayExpress repository include: E-MEXP-2032, E-MTAB-3309, E-MTAB-5095, E-MTAB-5913, and E-MEXP-3890. The scripts for performing the analyses are stored in the GitHub repository at https://github.com/ND91/PRJ0000004_MACMETA. The macIDR package is stored in the GitHub repository at https://github.com/ND91/macIDR.

## Supplementary Materials

### Supplemental methods

#### Microarray data preparation

All analyses were performed in the R statistical environment (v3.5.0). Download of the raw and processed GEO and AE microarray was performed using the GEOquery (v2.48.0) and the ArrayExpress (v1.40.0) packages respectively. Raw data was normalized in a platform-specific fashion: Affymetrix microarrays were normalized using the rma function from the affy (v1.54.0) and oligo (v1.40.2) packages, whereas Illumina and Agilent microarrays were normalized using the neqc function from the limma (v3.31.14) package. Quality control of the log_2_ transformed expression values was performed using WGCNA (v1.51) and arrayQualityMetrics (v3.32.0) to remove unmeasured samples, genes, and studies of insufficient quality. Microarray probes were reannotated to the Entrez ID according to the annotation files on Bioconductor. Probes that associated to multiple Entrez IDs were removed and multiple probes associating to the same Entrez ID were summarized by taking the median.

#### RNA sequencing data preparation

Public raw sequencing reads were sourced from the NCBI Sequence Read Archive (SRA) and converted to fastq files using the fastq-dump function from the SRA-tools package (v2.9.0). All raw fastq files were first checked for quality using FastQC (v0.11.7) and MultiQC (v1.4). The sequencing reads were aligned against the human genome GRCh38 using STAR (v2.5.4). Post-alignment processing was performed using SAMtools (v1.7) after which reads overlapping gene features obtained from Ensembl (v91) were counted using the featureCounts program in the Subread (v1.6.1) package. Gene annotations were converted from Ensembl IDs to Entrez IDs using biomaRt.

#### macIDR package

The macIDR package for R comprises a set of functions for performing macrophage classification analyses and visualization thereof using the log odds ratios learned from the aggregated datasets and can be downloaded from https://github.com/ND91/macIDR. Included in this package are the log odds ratios of the 500 repetitions and the logistic regression function necessary to perform classification. By default, the log odds ratios for each predictor gene were defined as described before, namely by taking the median of the log odds ratios for the stable genes across all 500 iterations. We acknowledge that some users might prefer different methods and have therefore provided the option for users to define their own stability threshold and method for aggregation when defining the input log odds ratios for classification analysis. The logistic regression approach implemented in macIDR ignores missing values as we reasoned that missing predictor genes should not prevent classification. Nonetheless, we provide functions such that the user can assess which genes and what relative proportion of log odds ratio is missing by calculating the relative log odds ratios. To that end, we leave it up to the user’s discretion to decide their own threshold for continuing with the classification analysis.

#### Functional analyses

Pairwise gene correlations were generated by calculating the Spearman correlation to minimize the effect of outliers. Pathway and regulator analyses were performed using the Canonical Pathways and Ingenuity Upstream Regulator found in the Ingenuity Pathway Analysis software package (QIAGEN Inc., https://www.qiagenbioinformatics.com/products/ingenuity-pathway-analysis) and Metascape (http://metascape.org/gp/index.html). Transcription factor motif analysis was performed by using HOMER (v4.10) using the following parameters: -start -200 -end 100 -len 8, 10, 12.

### Supplemental figures

**Fig. S1.**
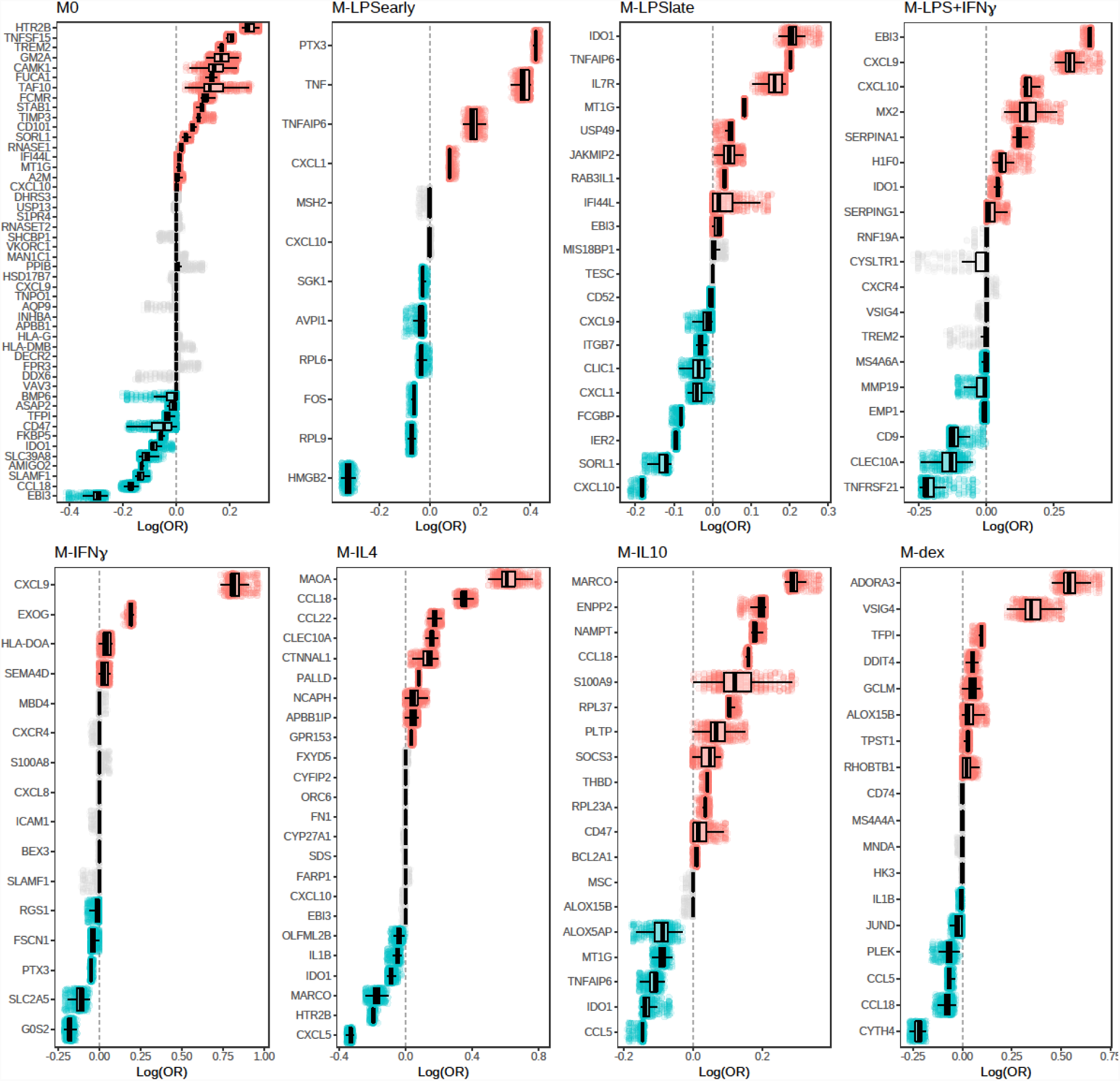
Median-stabilization. Plot representing the log odds ratio distribution as obtained from the 500 iterations of ten-fold cross-validation. Red and blue represent stable predictor genes with positive and negative coefficients respectively. Grey represents unstable predictor genes.

**Fig. S2.**
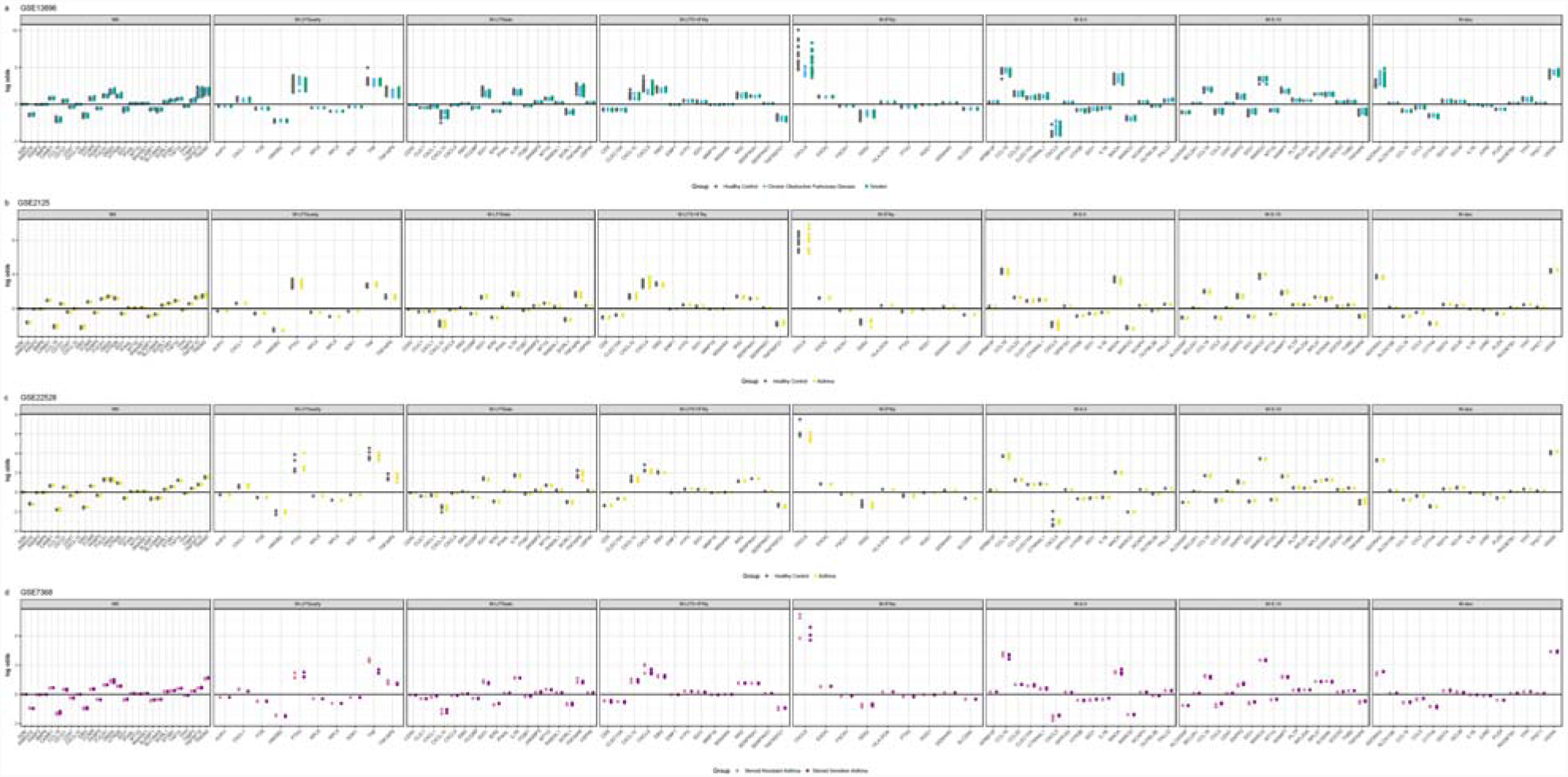
Log odds per gene for alveolar macrophage classification. Plots of the log odds per gene for the studies facetted by the different macrophage class for studies: (a) GSE13896, (b) GSE2125, (c) GSE22528, and (d) GSE7368.

**Fig. S3.**
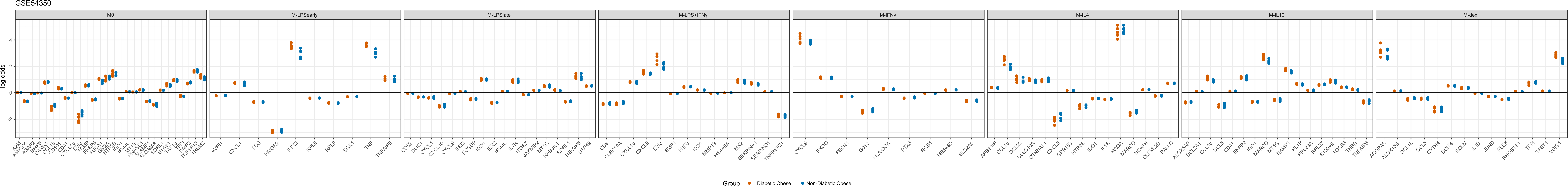
Log odds per gene for adipose tissue macrophage classification. Plots of the log odds per gene for the studies facetted by the different macrophage class for study: GSE54350.

**Fig. S4.**
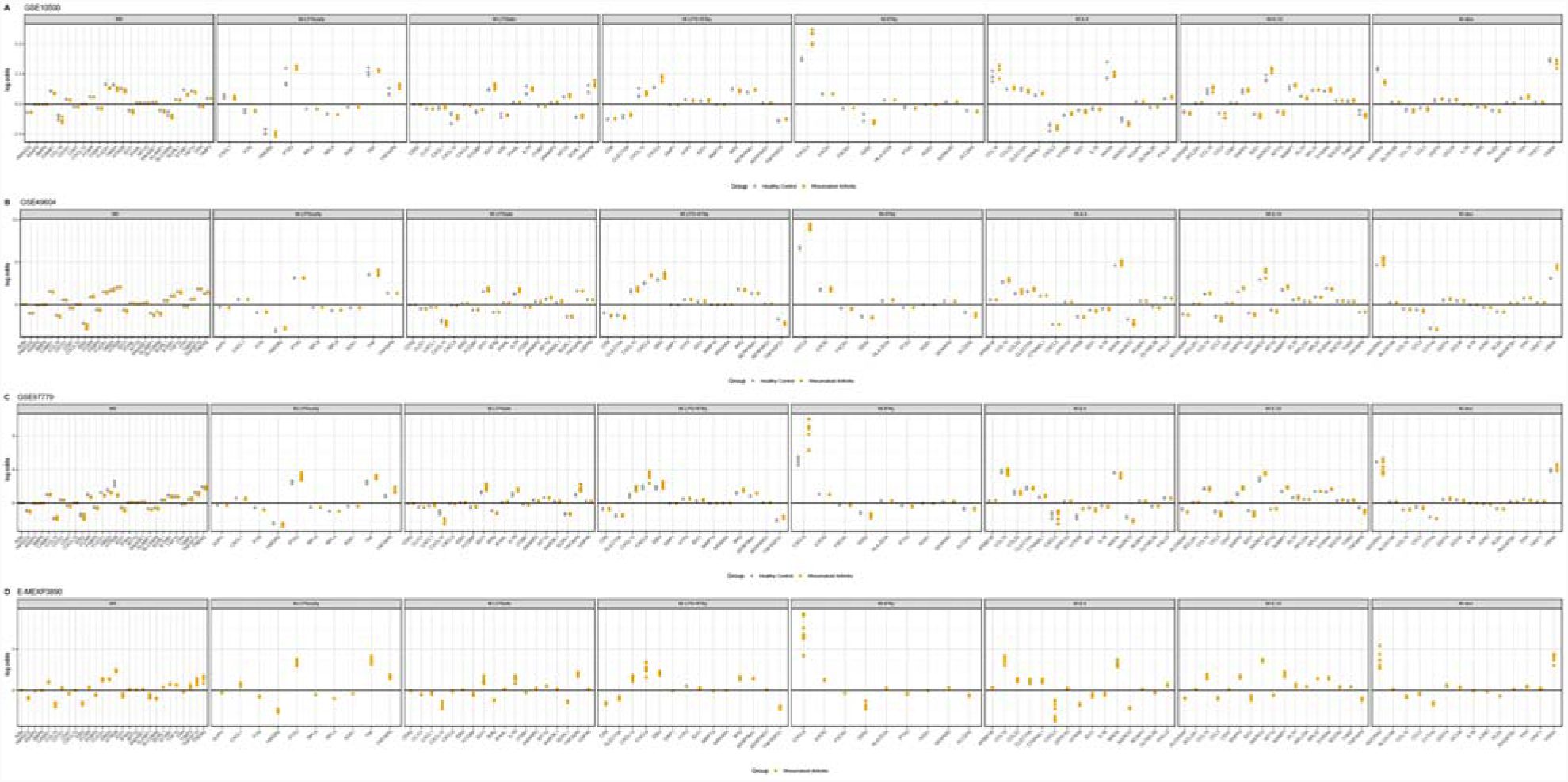
Log odds per gene for the synovial macrophage classification. Plots of the log odds per gene for the studies facetted by the different macrophage class for studies: (a) GSE10500, (b) GSE49604, (c) GSE97779, and (d) E-MEXP3890.

### Supplemental tables

**Table S1 meta-analysis results.** A table containing the unbiased estimator of the effect size and the variance as calculated by the meta-analysis ranked by the *p*-value. Separate tabs contain the results for the different comparisons of the meta-analysis. Columns “Entrez” and “Gene” represent the Entrez gene ID and the HGNC symbol respectively. The column “mu” represents the unbiased estimator of the effect size and the “mu_var” represents the unbiased estimator of the variance. The columns “Z”, “Z_pval”, and “Z_pval”BH” represent the Z-statistic, the associated *p*-value and the Benjamini-Hochberg adjusted *p*-value.

**Table S2 meta-analysis pathway analysis.** A table containing the aggregated results of the enriched canonical pathways for each comparison made for the meta-analysis.

**Table S3 predictor gene pathway analysis.** A table containing the aggregated results of the enriched canonical pathways from IPA (Tab 1) and pathway analysis from metascape (Tab 2) for all predictor genes.

**Table S4 Log odds for the verification datasets.** An Excel file where each tab represents the classification signal as depicted in log odds for each individual study. Each tab contains a table representing the log odds separated by predictor genes to depict the contribution of each gene to the classification. Each row represents a predictor gene for a particular class and each column represents a sample. Missing values for particular rows indicates that these genes were not present in the study. The last three columns represent the Entrez gene ID, HGNC gene symbol and the class to which the weights belong.

